# Size-dependent invasion and therapeutic phenotype of 42MGBA glioblastoma spheroids

**DOI:** 10.1101/2025.07.09.663980

**Authors:** Sheridan Fok, Anagha Shreesha, Angela Appiah-Kubi, Rebecca B. Riggins, Brendan A.C. Harley

**Author notes:** **Corresponding Author:** B.A.C. Harley, Dept. of Chemical and Biomolecular Engineering, Cancer Center at Illinois, Carl R. Woese Institute for Genomic Biology, University of Illinois at Urbana-Champaign, 110 Roger Adams Laboratory, 600 S. Mathews Ave., Urbana, IL 61801.

## Abstract

Glioblastoma (GBM) is one of the most common malignant brain tumors, with patient mortality driven by invasion into the surround brain microenvironment and drug resistance. Multicellular spheroids are increasingly a common model to study GBM invasion and drug response in engineered biomaterials. However, a key design feature of tumor spheroid studies is the size of each spheroid (number of cells, diameter). Given the heterogenous growth of GBM cells at the surgical margin, spheroids of different sizes may also have clinical relevance. Here, we define shifts in behavior and drug response of wild type and temozolomide (TMZ) resistant GBM spheroids as a function of initial spheroid size. GBM spheroids ranging from 100 to 12,000 cells in size were embedded into a methacrylamide-functionalized gelatin (GelMA) hydrogel. GBM spheroid size had an inverse relationship with the number of apoptotic cells. We observed significant spheroid size dependent effects on TMZ efficacy for both TMZ resistant and wild type cells. Interestingly, high single doses of TMZ were more effective in reducing three-dimensional migration from smaller spheroids than metronomic dosing while high single dose and metronomic dosing were equally effective in reducing invasion for large TMZ-resistant spheroids. Our study highlights the importance of considering and reporting spheroid size for cancer tissue engineering studies considering invasion and drug resistance. It also informs future studies of residual GBM at the tumor margins most responsible for patient relapse and mortality.

## 1. Introduction

Glioblastoma (GBM) is the most common, aggressive, and lethal form of primary brain cancer ^1,2^. While advances in radio- and chemotherapy have extended median survival from 12 to ∼15mo, overall survival (<5% after 5 yrs) remains poor and largely unchanged over the past two decades ^3-10^. Current standard-of-care includes maximal surgical resection followed by radiation and chemotherapy with the alkylating agent temozolomide (TMZ) ^11^. However, mortality is driven by rapid diffusive spreading of GBM cells beyond the surgical margin ^12-15^. GBM tumors recur rapidly (6.9mo post debulking) and in close proximity (>90% within 2cm) of the original tumor ^16,17^. It is therefore important to develop strategies to target GBM cells that spread diffusely into tissue microenvironment beyond the surgical margin ^18^. While studies of GBM progression and therapeutic response are commonly performed using *in vivo* assays ^19-22^, such models are less well-suited to examine both the kinetics of cell invasion as well as for nuanced studies of the role of the tumor microenvironment on progression.

*In vitro* experiments using both two-dimensional cell cultures ^23,24^ as well as three-dimensional biomaterial models of the brain tumor microenvironment ^15,25-27^ offer a lens to elucidate the instructive nature of the composition, mechanical properties, and multicellular nature of the GBM tumor microenvironment. Recent efforts have identified key features of the GBM microenvironment such as chemical and mechanical gradients ^28-30^ as well as fluid flow ^31-33^. Our group has developed a methacrylamide-functionalized gelatin (GelMA) hydrogel to study the role of matrix stiffness, hypoxia, and hyaluronic acid content on GBM progression using both immortalized cell lines and patient-derived xenografts ^34-36^. In these studies, metrics of GBM response can be gathered from low-density cell populations dispersed in, or high-density cell spheroids embedded into, the hydrogel network. Both modes of study are important because single cells and clusters of GBM cells may exist in the tumor margins after debulking ^37-40^. We used these hydrogel models to show angiocrine signals influence GBM invasion and TMZ resistance ^41-44^ and more recently to study the emergence of TMZ resistance ^45^ in response to physiologically-relevant single and metronomic ^46,47^ TMZ doses using isogenic pairs of TMZ-resistant and responsive GBM cell lines developed by *Tiek et al.* ^48^. Metronomic dosing offers an alternative to a higher single dosage ^46^, has been suggested to offer better penetration and potentially circumvent resistance mechanisms seen with high single drug doses ^49^, and is more consistent with the clinical use of TMZ. Our previously used GBM spheroids to profile patterns of cell invasion in response to changes in matrix composition, hypoxia, and the presence of GBM stem cell subpopulations ^36,50-53^. However, previous studies of the role of confinement and biophysical signals on liver development ^54-56^ and other cancer stem cell phenotypes ^57^ suggest spheroid size may be an essential design principal. Larger spheroids have been hypothesized to affect the emergence of hypoxia and necrosis as well as to induce changes in gene expression that may underlie invasion and chemoresistance ^58^. However, given the likelihood that multiple scales of GBM spheroids are present in the tumor margins it is essential to understand the degree spheroid size influences metrics of GBM phenotype.

The goal of this project is to define the influence of GBM spheroid size on cell activity, invasion, and drug response within a model GelMA hydrogel (Figure 1). We fabricated spheroids from both TMZ-resistant or wild type (WT) GBM cell lines, with sizes ranging between 100 and 12,000 cells per spheroid. We report the effect of spheroid size on GBM cell growth and gene expression, apoptosis, and invasion. We define the effect of single vs. metronomic TMZ dosing on GBM spheroids as a function of size and TMZ-resistance status. Together, these establish design principles regarding spheroid sizes necessary to evaluate therapeutic compounds as well as to establish biophysical thresholds (e.g., spheroid surface area to volume) that alter GBM phenotypes *in vitro*.

**Figure 1.**
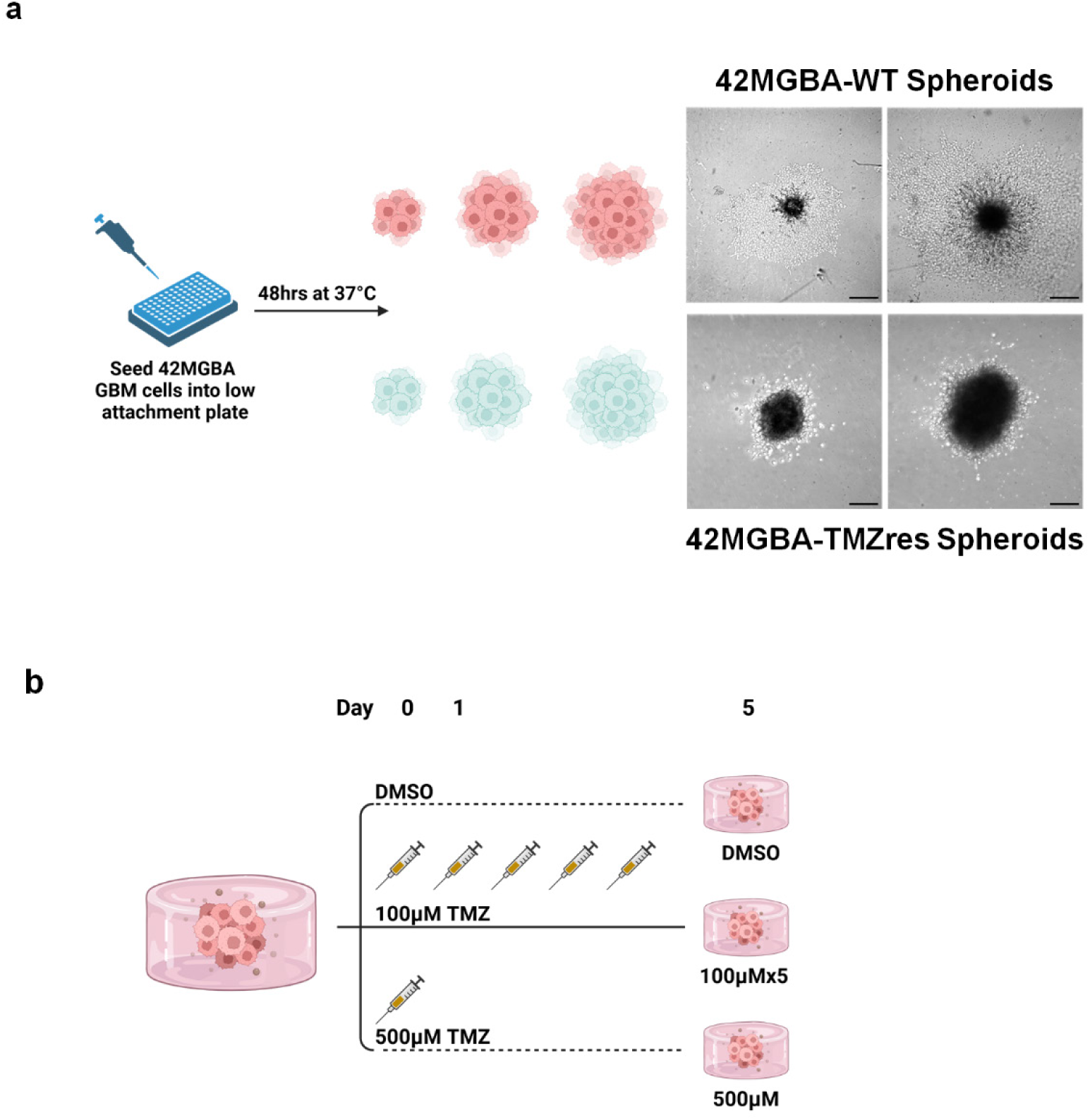
Conceptual illustration of experimental plan. (a) GBM spheroids were formed form isogenically-matched pairs of wild type (42MGBA-WT) or TMZ-resistant (42MGBA-TMZres) cells across a range of spheroid sizes (100 – 12,000 cells/spheroid). (b) Schematic depiction of the TMZ treatments. GBM spheroids encapsulated in GelMA hydrogel are subjected to a DMSO control, metronomic treatment (100µM per day for 5 days) via TMNZ, or single high TMZ dose (500µM at day 1). Cell phenotype, invasion, and gene expression patterns were traced for up to 5 days after initial TMZ exposure.

## 2. Materials and Methods

### 2.1. Cell culture

Wild type (42MGBA-WT) and Temozolomide resistant (42MGBA-TMZres) 42MGBA cell pairs were acquired from the Riggins lab at Georgetown University ^48^. 42MGBA cells were cultured using Dulbecco’s Modified Eagle Medium (DMEM) with 10% fetal bovine serum (FBS) and 1% penicillin/streptomycin; media was changed every 48 hours. Cells were maintained in incubators at 37°C and 5% CO_2_ and used within 10 passages.

### 2.2. Spheroid formation

GBM spheroids were generated using two well established protocols. <u>Ultra-low attachment plate method</u>: Wild type (42MGBA-WT) and TMZ resistant (42MGBA-TMZres) GBM cells were formed into spheroids using the nutation method ^44,59^. Briefly, GBM cells were seeded in Corning® spheroid microplates, incubated at 37*°C* on a rotating incubator shaker (ThermoFisher, Canoga Park, CA) at 60 rpm for 48 hours. <u>Aggrewell™ microwell method</u>: Wild type (42MGBA-WT) and TMZ resistant (42MGBA-TMZres) GBM spheroids were also generated at a lager quantity required for flow cytometry analysis. Briefly, AggreWell 400 or 800 microplates (STEMCELL Technologies, Vancouver, Canada) were pre-treated with anti-adherence rising solution (STEMCELL Technologies, Vancouver, Canada) for 5 minutes. The anti-adherence rinsing solution was then removed and replaced with 2 mL cell culture media seeded with the desired number of cells to form each spheroid.

### 2.3. Immunostaining

Sample were fixed using formaldehyde solution 4 % (Sigma Aldrich, Burlington, MA), permeabilized using 0.5% Tween-20(Fisher Scientific, Fairlawn, NJ) in PBS and blocked using 2% donkey serum (Sigma Aldrich) in PBS-T (0.1% Tween-20 in PBS) for 2 hours at room temperature. Ki-67 Recombinant rabbit monoclonal antibody SP6(ThermoFisher, Canoga Park, CA) was diluted at 1:200 ratio in 2% donkey serum in PBS-T. Samples were incubated in primary antibody at 4*°C* overnight. Alexa 555 was used as secondary antibody (ThermoFisher, Canoga Park, CA) at 1:500 ratio in PBS-T. Hydrogels were washed with PBS-T between antibody incubation. Hoescht 33342 (ThermoScientific, Rockford, IL) was used as a nuclear marker at a 1:1000 ratio in PBS-T.

### 2.4. Flow cytometry

All flow cytometry measurements were performed using a FACSymphony A1 configuration (BD Bioscience, Franklin Lake, NJ). Briefly, spheroids were recovered from biomaterial culture, dissociated using 0.5% Trypsin, then resuspended in 500 μL 5% FBS for analysis in biological triplicate. Live cells were gated by forward scatter (FSC) and side scatter (SSC). Fluorescence data was collected for GFP (excitation laser: 488nm; bandpass filter: 530/30nm) and PE-Texas Red (excitation laser: 561nm; bandpass filter: 610/20nm) with a minimum of 50,000 singlet events were collected for each sample. Flow cytometry data was analyzed using FACSDiva^TM^ Software (BD Bioscience, Franklin Lake, NJ) then graphed using Prism 8 software (GraphPad). Apoptotic cells were gated as PE-Texas Red (Bobo-3 iodide, Thermo Fisher Scientific, Waltham, MA)) and FITC (CalceinAM; Thermo Fisher Scientific, Waltham, MA) positive. Necrotic cells were gated as PE-Texas Red (Bobo-3 iodide, Thermo Fisher Scientific, Waltham, MA) positive and FITC (CalceinAM) negative. (Figure S2).

### 2.5. GFP transduction of 42MGBA Cell lines

42MGBA cells were transfected using the CMV-rFLuc-T2A-GFP-mPGK-Puro lenti-labeler^TM^ plasmid (Systems Bioscience, Palo Alto, CA) per manufacturer’s instructions. Briefly, 7,500 42MGBA cells were seeded in 48 well plates with both TransDux Max and Max enhancers diluted in culture media at a final concentration of 1x. The multiplicity of infection (MOI) used was calculated based on the infectious forming units (IFUs) provided by the manufacturer, with lenti-labeler added to the cells at an expected MOI of 50. Cells were incubated at 37°C and 5% CO_2_ for 72 hours before sorting (BigFoot Invitrogen ThermoFisher Spectral Cell Sorter, ThermoFisher, Canoga Park, CA) for GFP positive cells. Isolated GFP expressing cells were expanded for 2 passages before experiments.

### 2.6. GelMA synthesis

GelMA macromers for all hydrogel culture were synthesized as previously described ^60^. Briefly, porcine gelatin type A, 300 bloom (Sigma Aldrich, St. Louis, MO) was dissolved in carbon-bicarbonate buffer (pH 9.4) at 65 °C. 15 µL methacrylic anhydride (Sigma Aldrich, St. Louis, MO) was added dropwise per gram of gelatin, and the reaction proceeded for 2 hrs with continues stirring (250 RPM). The reaction was quenched with pre-heated deionized water and dialyzed for seven days against deionized water with daily exchange. The product was then frozen at -20°C and lyophilized for 7 days. HNMR was used to determine the degree of functionalization (DOF). GelMA with DOF of 85% (data not shown) was used in this study.

### 2.7. Hydrogel gel encapsulation and Invasion assay

GFP expressing 42MGBA-WT or 42MGBA-TMZres spheroids was suspended in a 5wt% GelMA precursor solution and then pipetted into 20 μL volume (cylindrical) Teflon molds (1 spheroid per hydrogel construct). GelMA hydrogels were polymerized in the presence of lithium acylphosphinate (LAP, 0.1% w/v) and exposed to UV light for 35s using a UV lamp (*λ* = 365 nm, 5.79 mW/cm^2^) ^61^. GBM cell invasion assay was performed using a DMi8 Yokogawa W1 spinning disk confocal microscope with a Hamamatsu EM-CCD digital camera (Leica Microsystems, Buffalo Grove, IL) per previously described methods ^44^. Overall GBM cell outgrowth area was quantified by using ImageJ (NIH, Bethesda, MD). Briefly, sequential slices (5 μm thick) were imaged along the hydrogels’ depth. Maximum intensity projections were assembled from approximately 50 slices, with the composite image then binarized and analyzed with the analyze particle function on ImageJ (NIH, Bethesda, MD). Six independent spheroid outgrowth measurements were gathered per experimental group as a biological replicate.

### 2.8. TMZ treatment

We performed TMZ studies in GelMA hydrogels as previously described ^45^. Briefly, temozolomide (TMZ, Calbiochem via Millipore Sigma, Burlington, MA) was dissolved in dimethyl sulfoxide (DMSO) at a concentration of 200 mM. DMSO alone (≦ 0.5% v/v) was used as a vehicle control for all experimental studies. Hydrogels containing GBM spheroids were cultured for 24 hours prior to the administration of TMZ. Hydrogels were subsequently treated with one of the following TMZ dose regimens: a single dose of 500 μM or five daily doses of 100μM (a cumulative dose of 500 μM). Media was replaced for both groups daily with either the appropriate concentration of TMZ or DMSO control.

### 2.9. Statistics

Statistics were performed using SPSS. Normality of data was determined using the Shapiro–Wilk test, and equality of variance was determined using Levene’s test. For normal data, comparisons between two groups were performed using a *t*-test, while comparisons between multiple groups were performed using a one-way ANOVA followed by Tukey’s post-hoc when assumptions were met. In the case where data was not normal or groups had unequal variance, comparisons between two groups were performed using a Mann–Whitney test, while comparisons between multiple groups were performed using a Kruskal–Wallis test with Dunn’s post-hoc. Significance was determined as *p* < 0.05. All quantitative analyses were performed on hydrogels or microfluidic assays set up across at least three independent experiments.

## 3. Results

### 3.1. Library of GBM spheroids as a function of cell size

We first created spheroids libraries for both wild type (42MGBA-WT) and TMZ-resistant (42MGBA-TMZres) GBM cell lines ranging in size from 100 cells to 12,000 cells (Figure 2A). We quantified the resultant size and morphology of the spheroids via overall projected surface area and circularity metrics from bright-field microscopy images. Spheroid projected surface area scale with numbers of cells and broadly, all spheroids displayed consistent circularity indices of greater than 60%. Spheroids generated from TMZ-resistant GBM cells were significantly larger than those generated from the same numbers of wild-type GBM cells (Figure 2B). And while not statistically significant, TMZ-resistant spheroids trended towards increased circularity than GBM wild type spheroids for all sizes up to up to 2500 cells; smaller (≤2500 cells) TMZ-resistant spheroids also trended toward higher circularity than TMZ-resistant spheroids of larger (≥5000 cells) size (Figure 2C). We characterized the packing of GBM cells using fluorescent images obtained from CMV-rFLuc-T2A-GFP-mPGK-Puro lenti-labeler^TM^ labeled wildtype and TMZ-resistant cells formed into 10,000 cell spheroids (Figure S1). TMZ-resistant spheroids appear to be more densely packed with a brighter GFP core, whereas the fluorescent signal for the wild type spheroids display a more diffuse organization.

**Figure 2.**
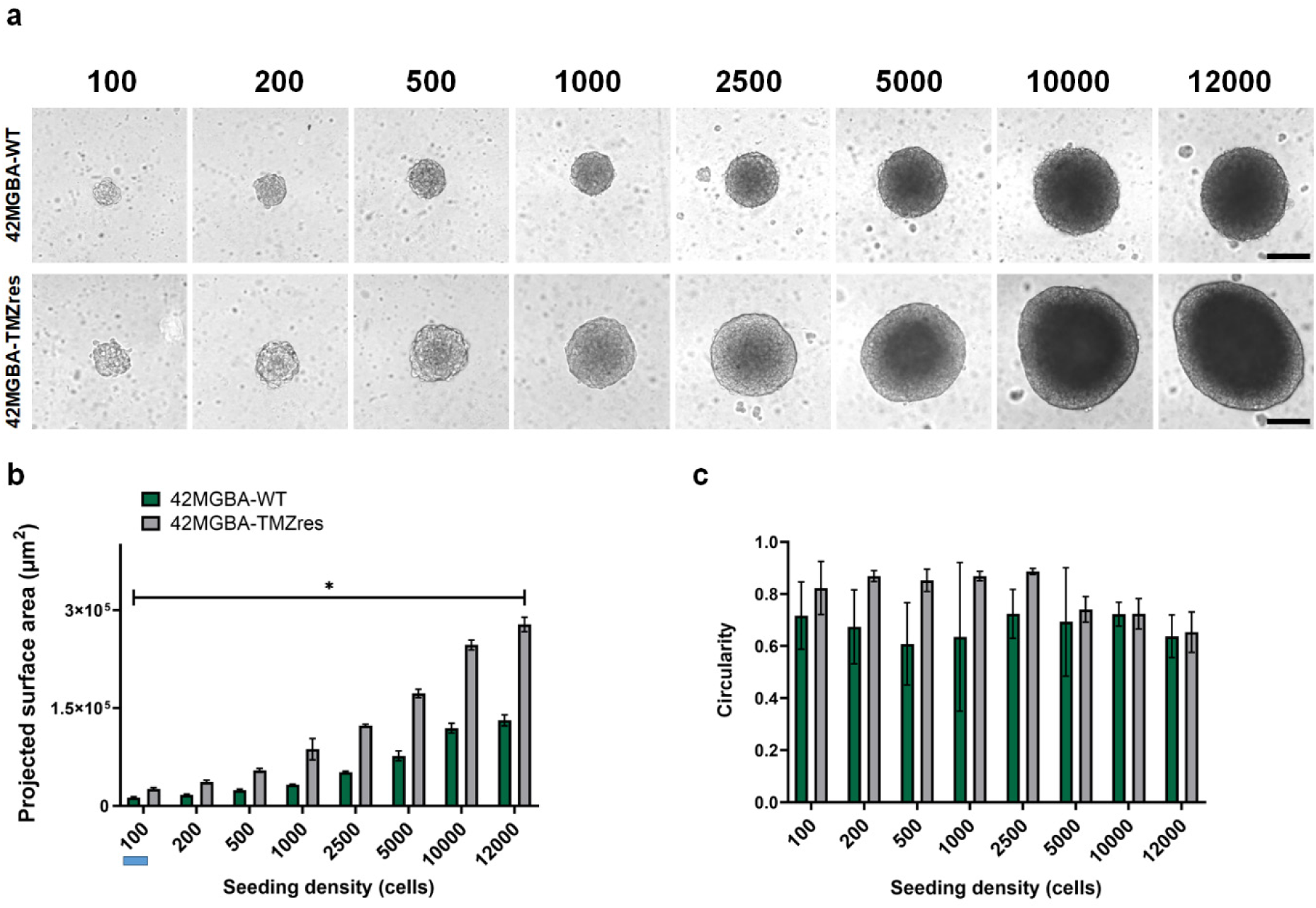
A library of GBM spheroids as a function of cell size. (a) Bright field image of 42-MGBA spheroids initiated with different number of cells after 48 hours (n=8/group). Scale bar: 200μm. (b) Projected surface area of GBM spheroids initiated as a function of size (n = 8/group). *: p < 0.05: statistical significance exists between WT and TMZres groups at each seeding density, as well as between each seeding density for WT and TMZres groups. (c) Circularity of 42MGBA spheroids initiated at different seeding densities (n= 8/group). Statistical analysis was done by Kruskal Wallis test followed by Mann Whitney U Test.

### 3.2. Patterns of GBM proliferation and apoptosis is influenced by spheroid size and TMZ-resistant status

We subsequently examined patterns of cell proliferation and death within GBM spheroids as a function of spheroid size and TMZ resistant status (42MGBA-WT vs. 42MGBA-TMZres). Immunostaining was performed 48 hours after encapsulation spheroids into GelMA hydrogels. Broadly, we observed Ki-67+ proliferative cells in all 42MGBA-WT spheroids regardless of spheroid size (Figure 3A). We observed higher levels of Ki-67 staining in 42MGBA-TMZres spheroids increasingly at the spheroid periphery, with the effect most pronounced in larger spheroids (Figure 3B). We also interrogated patterns of cell death in GBM spheroids as a function of spheroid size and TMZ-resistance status 48 hours after formation (Figure 3C, D). Broadly, bobo-3 iodide positive (dead) GBM cells were observed in spheroids of all sizes for both 42MGBA-WT and42MGBA-TMZres cells (Figure 3C, D)

**Figure 3.**
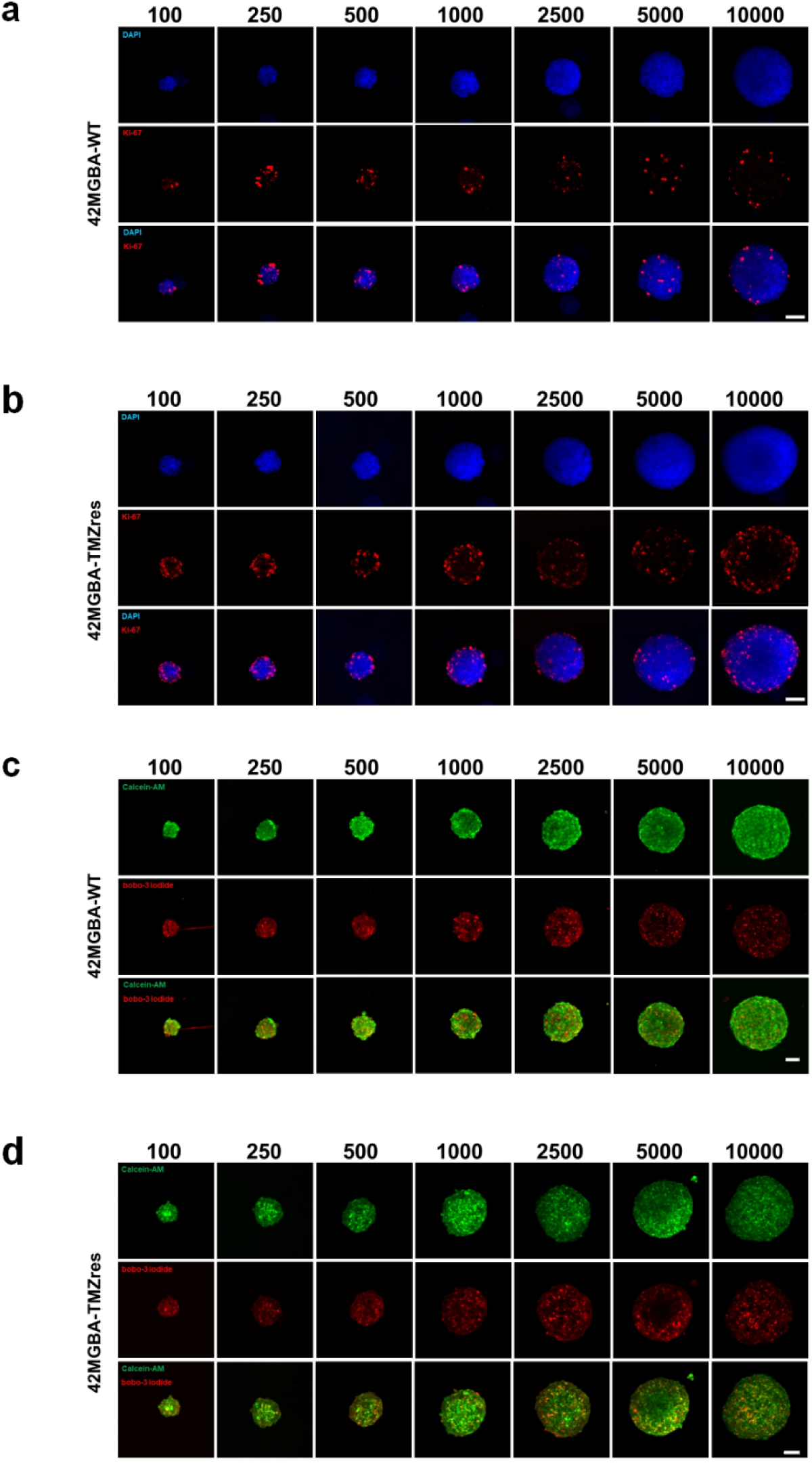
Patterns of GBM cell proliferation and apoptosis. Representative fluorescent images of (a) wild type vs. (b) TMZ-resistant spheroids labeled with DAPI (nuclei) and Ki-67 (G1/S/G2/M phases of the cell cycle) as a function of spheroid size (n=8/group). Representative fluorescent images of (c) wild type vs. (d) TMZ-resistant spheroids labeled with Calcein-AM (live cells) and bobo-3 iodide (dead cells) as a function of spheroid size (n= 8/group). Scale bar: 200μm.

To better assess whether spheroid size influenced the emergence of an apoptotic or necrotic GBM population, large quantities of spheroids of each size were generated using an AggreWell^TM^ microwell plate process. These spheroids were pooled by size then dissociated for flow cytometry analysis of the cellular content (Figure S2). Dissociation with 0.5% trypsin showed no effects on the GBM cell viability (data not shown). Disaggregated cells were analyzed for expression of bobo-3 iodide (PE-TexasRedA) and CalceinAM (FITC-A), with cells characterized as apoptotic (bobo-3+/CalceinAM+) vs. necrotic (bobo-3+/CalceinAM-). We observed few (<5%) overall necrotic cells and no significant difference in the fraction of necrotic cells in spheroids as a function of spheroid size or TMZ-resistance phenotype (Figure 4A).

**Figure 4.**
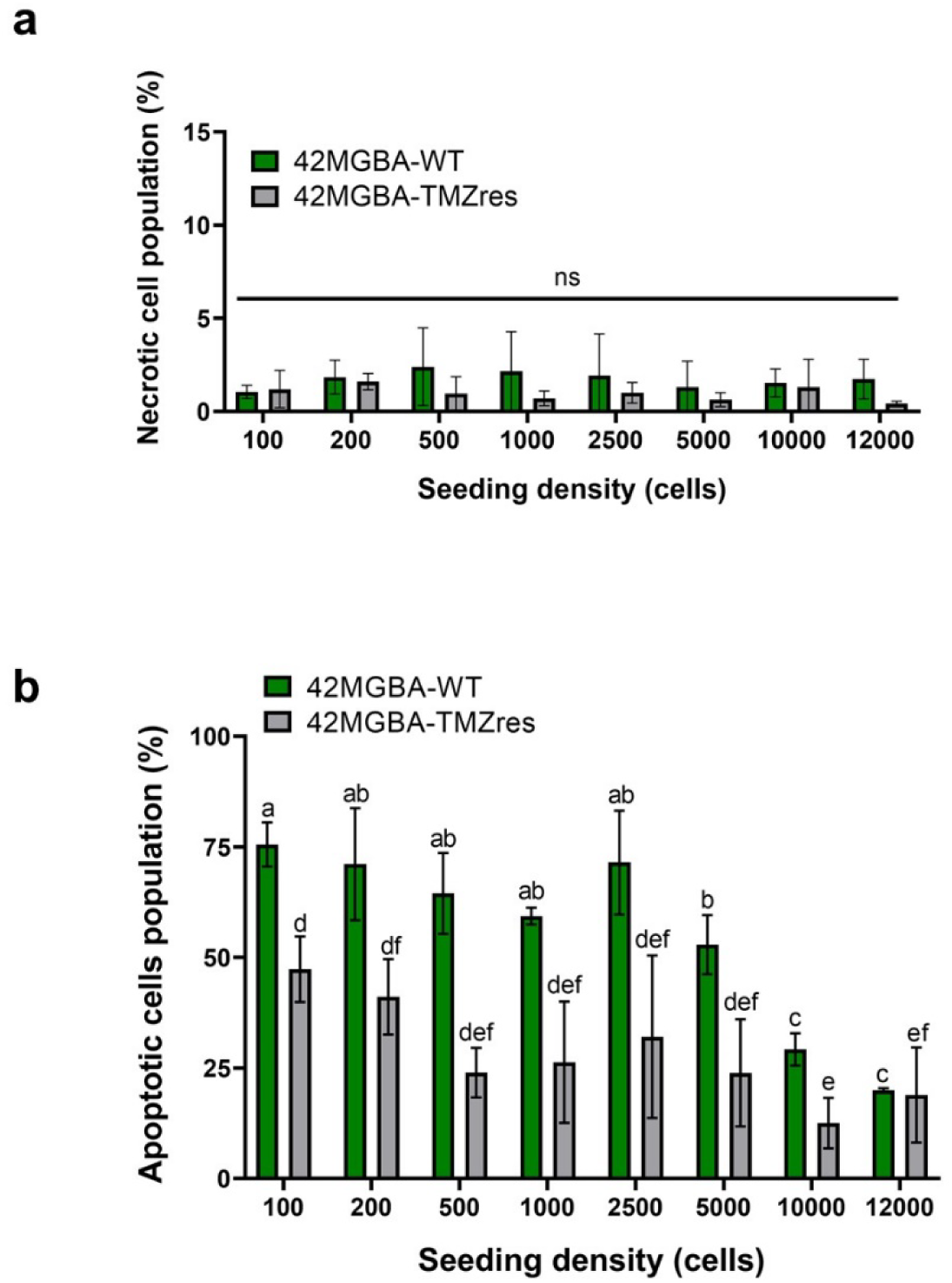
Percentage of apoptotic vs. necrotic GBM cells as function of spheroid size and TMZ-resistance. (a) The percentage of necrotic cells in spheroids as a function of spheroid size and TMZ resistance status (n=3). (b) The percentage of apoptotic cell population in spheroids as a function of spheroid size and TMZ resistance status (n=3). Statistical analysis was completed by One-way ANOVA followed by Tukey HSD Test.. Different letter (a, b, c) denotes p < 0.05 between wild-type spheroids as a function of size; different letters (d, e, f) denotes p <0.05 between TMZ resistant spheroids as a function of size.

Interestingly, we observed significant effects of both spheroid size and TMZ-resistance on the emergence of apoptotic GBM cells (Figure 4B). We observed significantly higher fractions of apoptotic cells in 42MGBA-WT vs. 42MGBA-TMZres spheroids for almost all spheroid sizes with higher fractions of apoptotic cells in smaller (≤ 5000 cells) than larger (10,000, 12,000 cells) spheroids regardless of TMZ-resistance status. For all subsequent studies we chose two spheroid condition (1,000 vs. 10,000 cells) that displayed different levels of proliferative capacity and apoptosis.

### 3.3 Single high dose and metronomic TMZ reduces GBM invasion for WT, but not TMZ-resistant spheroids

Spheroids initiated with 1,000 and 10,000 cells were encapsulated in GelMA hydrogel and subjected to treatment via TMZ. TMZ exposure was either a single exposure to 500 μM TMZ, replaced after 24 hours with complete medium containing equivalent concentration of DMSO, or metronomic dosing via 100 μM TMZ each day for 5 consecutive days, with all treatments compared to a DMSO control (Figure S3). Small (1,000 cells/spheroid) and large (10,000 cells/spheroid) 42MGBA-WT spheroids displayed significant invasion into the surrounding GelMA hydrogel, marked by increased in projected surface area with time. While trends emerged within 3 days of treatment, the effect of a single high dose or metronomic dosing on small 42MGBA-WT spheroids did not become significant until 5 days after treatment, with significantly reduced invasion in response to metronomic doing (vs. control) and a single high TMZ dose (vs. metronomic, control; Figure 5A, C). The effects of TMZ treatment on larger (10,000 cells/spheroid) GBM spheroids followed similar trends, though with a single high TMZ dose significantly reducing cell invasion as early as 1 day post treatment, and with metronomic dosing or a single high dose both significantly reducing invasion by day 5 (Figure 5B, D).

**Figure 5.**
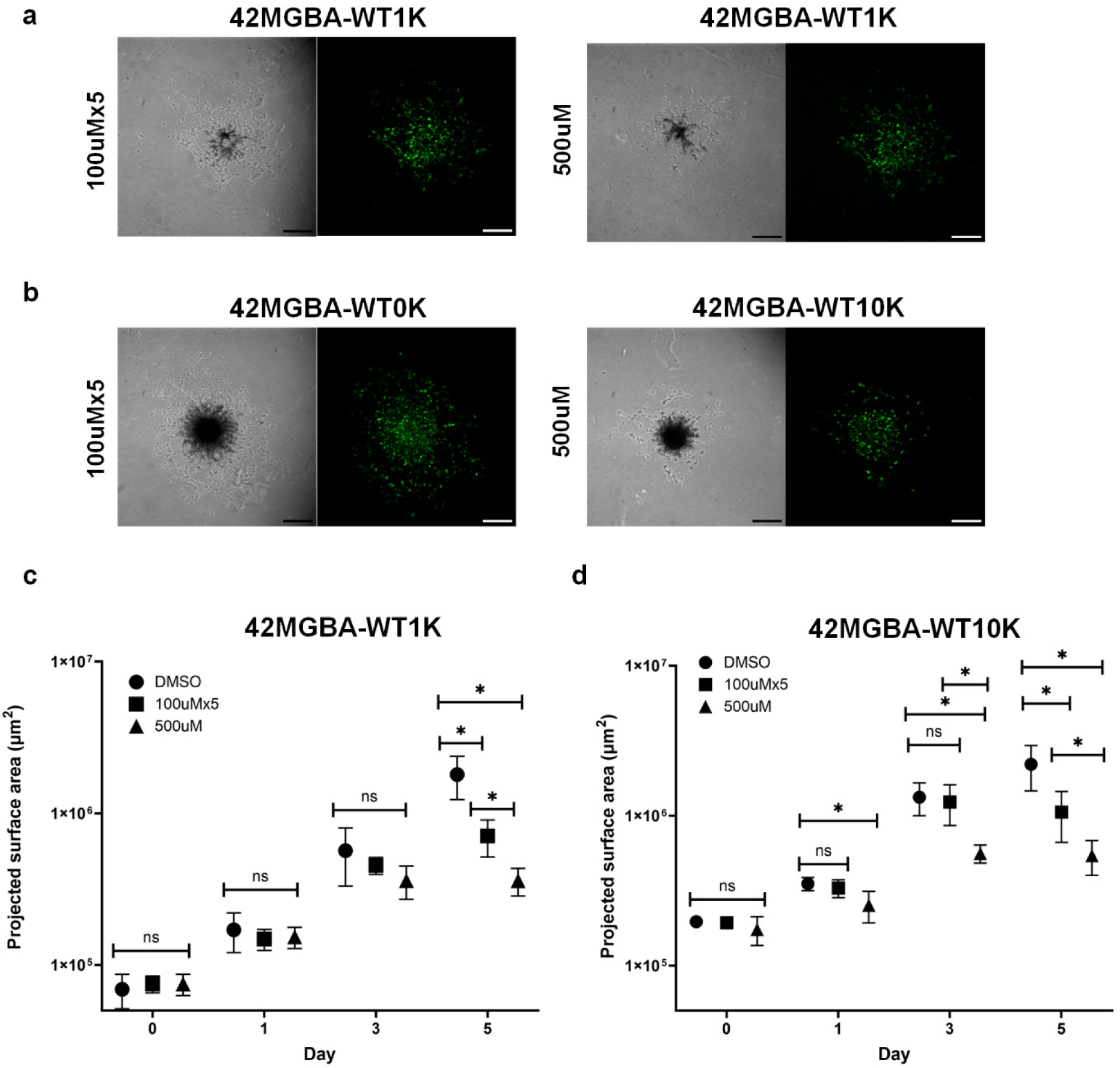
The effect of a single high TMZ dose or metronomic dosing on invasion of WT spheroids. (a) Representation image of small (1,000 cells) 42MGBA-WT spheroids treated with metronomic (left) vs. a single high TMZ dose (right). (b) Representation image of large (10,000 cells) 42MGBA-WT spheroids initiated with 10,000 cells treated with metronomic (left) vs. a single high TMZ dose (right). (c) Quantifying cell outgrowth for small (1,000 cell) spheroids over the course of 5 days after TMZ treatment (n = 3/group). (d) Quantifying cell outgrowth for large (10,000 cell) spheroids over the course of 5 days after TMZ treatment (n = 3/group). Statistical significance is represented as * p <0.05.

We observed limited invasion of 42MGBA-TMZres GBM cells for both small and large spheroids, consistent with earlier studies of 42MGBA-TMZres invasion in 3D hydrogels ^45,62^. Smaller spheroids (initiated with 1,000 cells) did not show any changes in cell invasion in response to either a single high dose or metronomic dosing (Figure 6A, C). While both TMZ treatment groups significantly reduced projected surface area on day 3 and 5 for large spheroids (Figure 6B, D), there was no different between TMZ treatments and overall invasion was small compared to wild type GBM cells.

**Figure 6.**
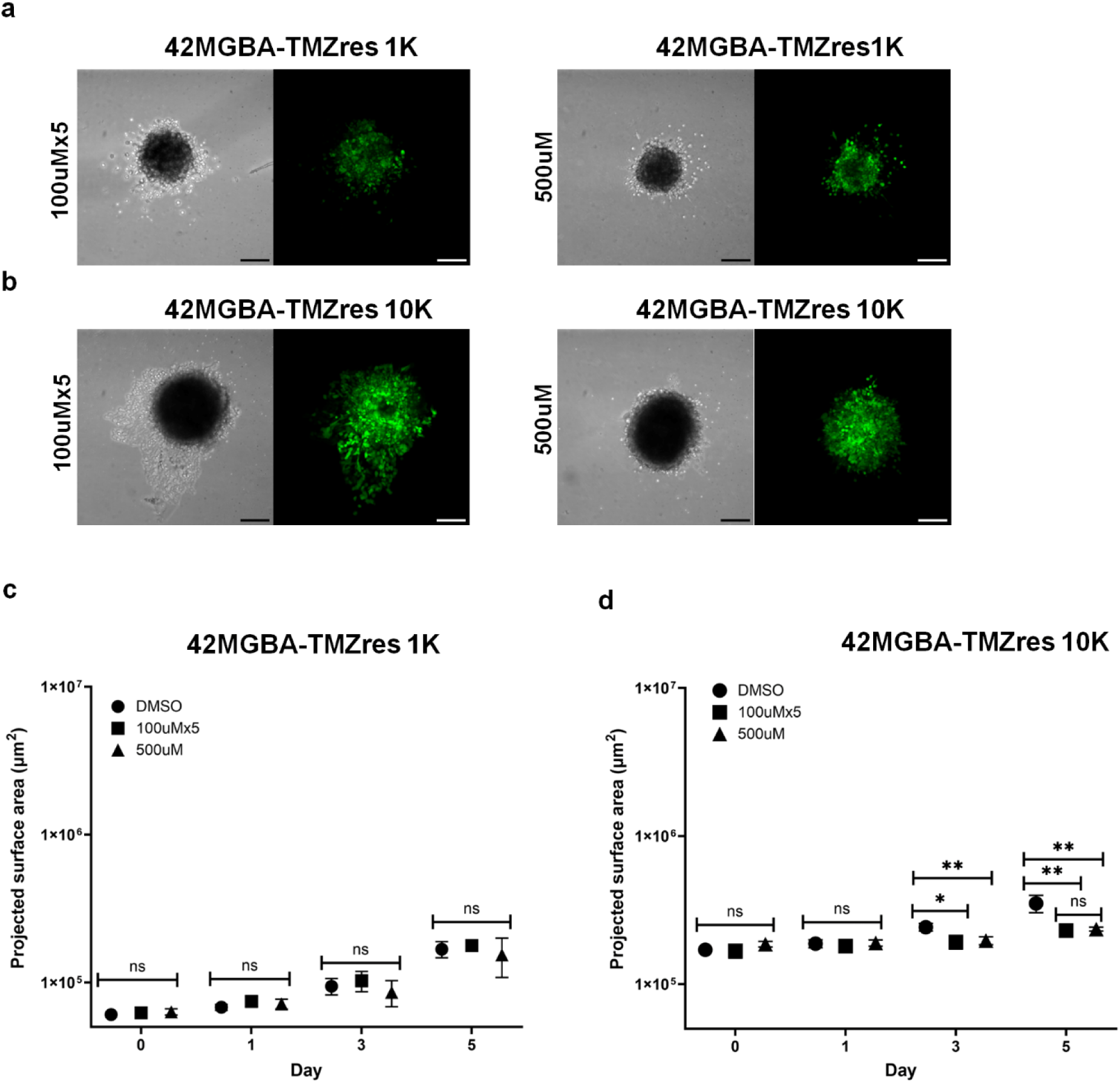
The effect of a single high TMZ dose or metronomic dosing on invasion of TMZ-resistant spheroids. (a) Representation image of small (1,000 cells) 42MGBA-TMZres spheroids treated with metronomic (left) vs. a single high TMZ dose (right). (b) Representation image of large (10,000 cells) 42MGBA-TMZres spheroids initiated with 10,000 cells treated with metronomic (left) vs. a single high TMZ dose (right). (c) Quantifying cell outgrowth for small (1,000 cell) TMZ-resistant spheroids over the course of 5 days after TMZ treatment (n = 3/group). (d) Quantifying cell outgrowth for large (10,000 cell) TMZ-resistant spheroids over the course of 5 days after TMZ treatment (n = 3/group). Statistical significance is represented as * p <0.05.

### 3.4. Difference in gene expression across spheroids of different sizes

We performed qRT-qPCR on gene targets related to cell death (*CASP3*, *TGF-β*), cellular growth (*PTEN*, *TIMP-1*), invasion (*MMP2*), and drug resistance (*CD44*, *HIF1α*) (Figure 7A). Wild type GBM cells in larger spheroids showed increased expression of *PTEN*, *STAT3*, *CASP3*, *TGF-β*, *TIMP-1* and *MMP2*, but reduced CD44 expression. Interestingly, a different expression pattern was observed for TMZ resistant cells, where larger spheroids demonstrated increased expression of *TIMP-1* and *CD44*, but reduced expression of *TGF-β*. Interestingly, quantitative analysis of *TIMP-1* (ΔCT normalized to actin expression) showed trends towards higher *TIMP-1* expression in larger vs. smaller spheroids for both wild type and TMZ resistant cells, but with TMZ resistant cells displaying significantly higher *TIMP-1* expression vs. wild type cells (Figure 7B).

**Figure 7.**
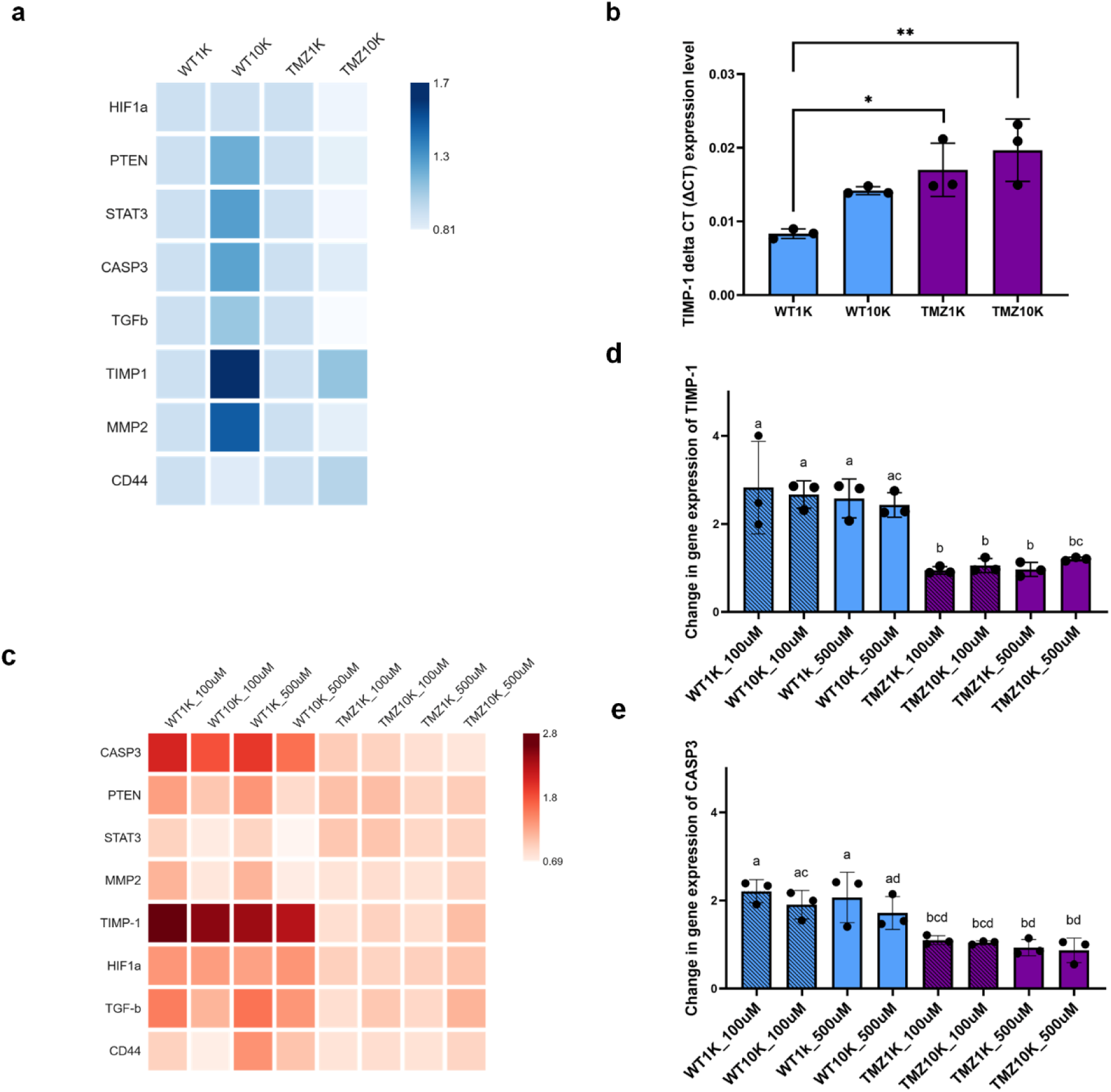
Shift in gene expression as a function of GBM spheroid size, TMZ treatment, and TMZ-resistance status. **Key: WT1k:** 42MGBA-WT, 1,000 cell spheroid; **WT10k:** 42MGBA-WT, 10,000 cell spheroids. **TMZ1k:** 42MGBA-TMZres, 1,000 cell spheroids; **TMZ10k:** 42MGBA-TMZ, 10,000 cell spheroids. **_100µM**: treated with 5 daily (metronomic) doses of 100µM TMZ. **_500µM**: treated with 5 metronomic doses of 100µM TMZ. (a) Heatmap of relative gene expression of wild type vs. TMZ resistance spheroids of different sizes (1000 or 10000 cells) in conventional culture. WT1K and TMZ1K are set to 1.0 for foldchange comparisons. (b) ΔCT value of *TIMP-1* of wildtype and TMZ-resistant spheroids of different sizes (1,000 vs. 10,000 cells). All data were normalized to ΔCT value of house-keeping gene Actin (n = 3/group). (c) Heatmap of relative gene expression of wild type vs. TMZ resistant spheroids of different sizes (1,000 vs. 10,000 cells) and as a function of TMZ treatment relative to DMSO control (single 500 μM dose vs. 5 days of 100 μM metronomic treatment). (d) *TIMP-1* expression for GBM spheroids as a function of size, TMZ resistance status and TMZ dose, relative to the DMSO control (n = 3/group). (e) *CASP3* gene expression for GBM spheroids as a function of size, TMZ resistance status and TMZ dose, relative to DMSO control (n = 3/group). Statistical analysis was completed by One-way ANOVA followed by Tukey HSD Test. Different letters denoted significance level at p < 0.05.

### 3.5. Change in gene expression on Day 5 after TMZ treatments

We subsequently examined changes in gene expression of GBM spheroids of different sizes after 5 days of metronomic or single high TMZ treatment. In general, wild type cells showed an increased expression (vs. DMSO control) of *CASP3*, *TIMP-1*, *HIF1α*, and *TGF-β* in response to either TMZ treatment (Figure 7C). *TIMP-1* expression was significantly higher in the wild type cell line compared to the TMZ resistance cell line after TMZ treatment (Figure 7D); while less pronounced, similar trends were also observed for increased *CASP3* expression in wildtype GBM cells after TMZ treatment (Figure 7E). Interestingly, small wild type spheroids showed higher expression of *PTEN*, *STAT3*, *MMP2*, *TGF-β* and *CD44* compared to the larger spheroids when treated with either metronomic or single high TMZ doses. Even though there was no statistical significance, several trends were observed in changes in gene expression for the TMZ resistant cells. Notably, metronomic dosing increased expression of *CASP3*, *PTEN*, and *STAT3* compared to a single high TMZ dose, while larger spheroids trended towards increased expression of *MMP2*, *TIMP-1*, *TGF-β*, and *CD44* in response to single high TMZ doses (Figure 7C, S4).

## 4. Discussion

Glioblastoma is marked by significant invasion into the surrounding brain parenchyma. Cancer tissue engineering platforms offer the opportunity to dissect the role of multiple components of the tumor microenvironment on metrics of progression and therapeutic response. Many first generation studies have used GBM cells distributed within a hydrogel biomaterial, simulating low-density cancer cell populations in order to assess how matrix composition ^50,63,64^, stiffness ^50,65-68^, and structure ^69^ influence invasion and drug response. While important, the multicellular nature of GBM clusters *in vivo* may also significantly influence cell activity. Yet while multicellular tumor spheroids are increasingly used in a wide range of cancer tissue engineering studies, spheroids sizes can differ drastically among studies ^70,71^. Larger spheroids are often hypothesized to lead to greater diffusional gradients for nutrients and oxygen that may contribute to the formation of necrotic core, regional changes in cell proliferative activity that may promote chemoresistance, and shifts in gene expression associated with a more aggressive phenotype ^72,73^.

The goal of this study was to define the role of spheroid scale on metrics of GBM progression (invasion, drug response) traditionally used in cancer engineering models. Because TMZ response can be tied to the presence of proliferative cells, tissue engineering models of TMZ response have often relied on supraphysiological TMZ doses. We recently validated a tissue engineering workflow, using an isogenically-matched pair of TMZ resistant and responsive cells with disparate MGMT expression levels generated by Tiek et al. ^48^, to assess TMZ response at physiological and metronomic dosing levels. While primarily for cells suspended in a GelMA hydrogel matrix, these studies confirmed that 42MGBA-TMZres cells showed reduced TMZ sensitivity in GelMA hydrogels. Our current study adds to this narrative, showing spheroid size is an important design characteristic that influence GBM cell behavior.

While we described spheroids from a fabrication perspective, it is notable that we created GBM spheroids from both 42MGBA-WT and 42MGBA-TMZres cells that range in scale from <130 µm (wild type 100 cells/spheroid) to >600 µm (TMZ-resistant 12,000 cells/spheroid) in diameter that are relevant for surgical resection and recurrence perspectives. While spheroid size increased with number of cells per spheroid and showed a high degree of circularity, spheroids created from TMZ-resistant cells were significantly larger, more compact, and trended towards a higher degree of circularity than wild type GBM cells. While qualitative fluorescent images suggested that Ki-67+ (proliferative) and bobo3+ (dead) cells were both more likely to be found towards the periphery of WT and TMZ-resistant spheroids, flow cytometry analysis of spheroids showed that a higher fraction of apoptotic cells were present in wild type (vs. TMZ-resistant) cells, particularly in smaller (<5000 cells/spheroid) spheroids. While previous work using mammary carcinoma cells ^76^ has suggested the formation of a necrotic core in tumor spheroids, we observed only a small fraction of necrotic cells. However, the increased fraction of apoptotic cells we observed in small GBM spheroids suggests that a relationship between spheroid size and apoptosis and necrosis may be more variable between cell types. There is an opportunity to explore extended time of spheroid culture to observe whether increased fractions of necrotic GBM cells can be observed. However, the increase in Ki-67 staining observed in GBM spheroids, particularly in larger spheroids containing TMZ-resistant GBM cells, was also consistent with increased 42MGBA-TMZres proliferation seen previously in 2D culture ^48^.

GBM invasion vs. proliferation (go vs. grow) plasticity suggests a process by which GBM cells can navigate complex selection pressures in the tumor margins ^77,78^. Interestingly, we observed that while 42MGBA-TMZres proliferation was higher in spheroid culture, both large and small spheroids formed from 42MGBA-WT cells displayed significantly increased invasion into GelMA hydrogels. This finding was also consistent with our prior work that showed for 5000 cell spheroids 42MGBA-WT cells were significantly more invasive than TMZ-resistant cells ^45^.

Importantly, our work here extends the range over which this observation is consistent (1000 – 10,000 cells/spheroid) and suggesting an invasive phenotype is exhibited by 42MGBA-WT cells across a wide range of biophysical environments. More interestingly, invasion of 42MGBA-WT cells was significantly reduced by both metronomic and a single high dose of TMZ, though the effect took 3 – 5 days to emerge. Interestingly, 42MGBA-TMZres_r_ invasion was largely insensitive to TMZ treatment except for large (10,000 cells/spheroid) spheroid cultures. Our prior work showed 42MGBA-TMZres cells (versus 42MGBA-WT) displayed reduced G2/M and increased G1 phase in 2D culture ^48^, and no change in metabolic activity in response to 100 µm TMZ in 3D GelMA culture^45^. Our current findings now suggest 42MGBA-WT cells also show reduced invasion in response to TMZ as well as the emergence of spatial patterns of cell proliferation vs. invasion in large 42MGBA-TMZres spheroids that may be consistent with go-vs. grow plasticity. These findings are different from prior work with ovarian cancer cells that showed larger (5000 cells) spheroids were more resistant to taxol and cisplatin compared to smaller (500, 200, 100 or 50 cells) spheroids ^79^. As larger spheroids have also been suggested to exhibited poor drug biotransport ^72^, our finding suggest the need for future study to more closely examine the spatial dependent shifts in GBM cell behavior and drug response. The observed differences in patterns of cell proliferation and spheroid-size dependent effects on apoptotic fraction, as well as the relative dearth of publications carefully interrogating the role of spheroid size in engineered cancer models, suggest the need for future studies that quantify radial patterns of cell proliferation, apoptosis induction, and metronomic TMZ exposure as a function of spheroid size and time.

Finally, our analysis of shifts in GBM gene expression suggested the largest changes were due to differences between TMZ-resistant and wild type cells, rather an effect of spheroid size. However, interestingly, TMZ-resistant (vs. wild type) cells displayed significantly upregulated expression of the MMP inhibitor TIMP-1. TIMP-1 has a multitude of functions, and has been implicated in suppressing cell proliferation and metabolic activity in mesenchymal stem cells ^80^, but increasing cell proliferation and invasion via the TIMP1/FAK/Akt pathway in colorectal cancer cells ^81^. Here, we observed increased TIMP-1 expression in large (10,000 cells) vs. small (1,000 cells) spheroids for both wild type and TMZ-resistant lines, and increased TIMP-1 expression in TMZ-resistant vs. wild type spheroids. We also observed phenotypic shifts in expression of Caspase-3 (*CASP3*), a protease implicated in programmed cell death. *CASP3* expression was elevated in wild type spheroids after both metronomic and single high TMZ doses, especially in small spheroids. Interestingly, while we did not see a difference in metronomic vs. a single high TMZ dose on invasion in TMZ resistant cells, the combination of metronomic dosing and small spheroids induced the greatest increase in *CASP3* expression in TMZ resistant cells (Figure 7E). We observed an inverse effect of spheroid size on *MMP-2* expression in 42MGBA-WT specimens as well as a modest increase with spheroid size for 42MGBA-TMZres specimens. These findings, while broadly consistent with prior studies using drug resistant (vs. wild type) glioblastoma cell lines that did not considering spheroid size ^48,82^, suggest that smaller GBM spheroids may display the most robust responses to TMZ treatment in tissue engineering models. This again argues more careful consideration of spheroid size as a critical design factor in tissue engineered cancer models.

## 5. Conclusions

We report the adaptation of a well-characterized GelMA hydrogel system to evaluate the combined effect of TMZ dosing strategy (metronomic vs. single high TMZ dose), TMZ-resistance status, and the size of GBM spheroids on metrics of GBM invasion and drug response. While spheroids embedded into a hydrogel matrix are increasingly common in studies of GBM cell behavior, an explicit consideration of the role of spheroid scale is essential to interpret these findings. TMZ-resistant and wild type GBM cells maintain their TMZ response phenotypes, regardless of spheroid size. Analysis of the spatial pattern of cell proliferation as well as underlying gene expression data suggests that smaller GBM spheroids may exhibit an increased fraction of apoptotic cells as well as a more robust response to TMZ treatment. These findings suggest that spheroid size should be rigorously reported and that future investigation must also consider the role of spheroid size on matrix remodeling (TIMP1) and apoptosis-associated (*CASP3*) proteases.

## Acknowledgements

Funding sources include the National Cancer Institutes of the National Institutes of Health under Awards Number R01 CA256481 (BACH), R01 CA279195 (R.B.R.), and T32 CA009686 (fellowship funding for A.A-K.; T32 principal investigator (PI): Dr. Chunling Yi). The content is solely the responsibility of the authors and does not necessarily represent the official views of the NIH. The authors are also grateful for additional funding provided by the Department of Chemical & Biomolecular Engineering and the Cancer Center at Illinois at the University of Illinois Urbana-Champaign. The authors also wish to acknowledge assistance from the Core Facilities at the Carl R. Woese Institute for Genomic Biology as well as Dr. Marcin Wozniak at the Cytometry and Microscopy to Omics Facilities at the Roy J. Carver Biotechnology Center for training and assistance with Flow Cytometry. Figures were created in BioRender.com.

## Contributions (CRediT: Contributor Roles Taxonomy^83,84^)

**S. Fok:** Conceptualization, Data curation, Formal Analysis, Visualization, Investigation, Methodology, Writing – original draft, Writing – review & editing. **A. Shreesha:** Investigation. **A. Appiah-Kubi**: Resources, Writing – review & editing. **R.B. Riggins:** Resources, Funding acquisition, Writing – review & editing. **B.A.C. Harley:** Conceptualization, Resources, Project administration, Funding acquisition, Supervision, Writing – review & editing.

