## Supplemental Information for "Size-dependent invasion and therapeutic phenotype of 42MGBA glioblastoma spheroids"

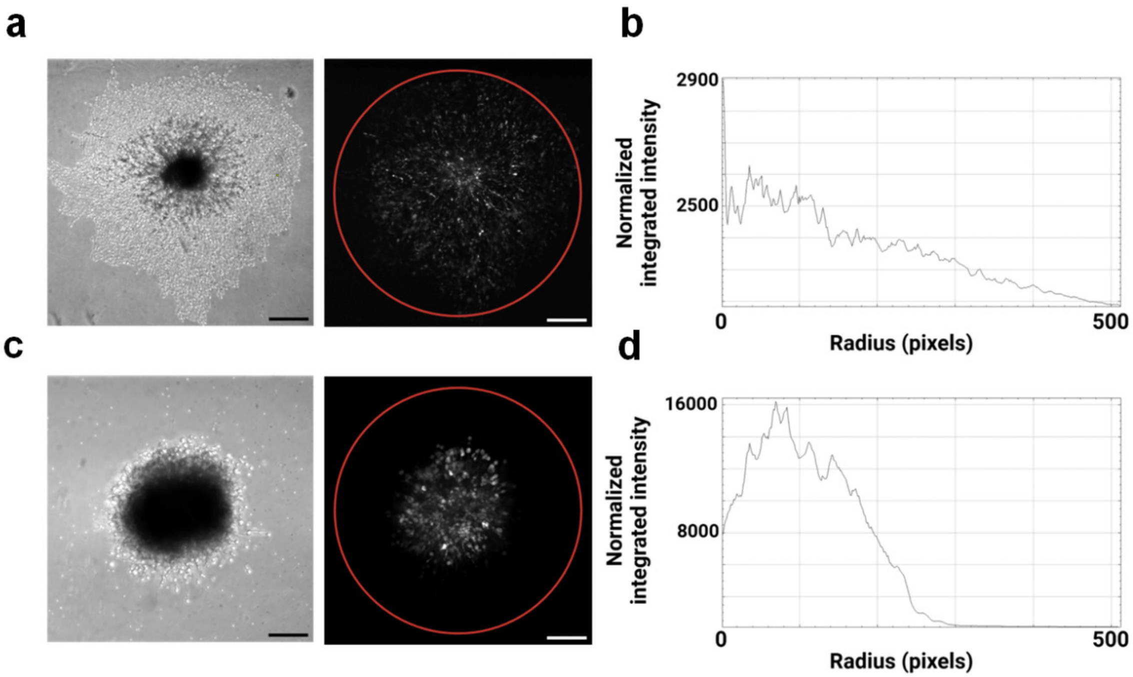


**Figure S1.** (a) Representative image of 42MGBA-WT spheroid initiated with 10,000 cells; bright field (left), grayscale of the GFP channel (right). (b) radial analysis of the fluorescent intensity. (c) Representative image of 42MGBA-TMZres spheroid initiated with 10,000 cells; bright field (left), grayscale of the GFP channel (right). (d) radial analysis of the fluorescent intensity. Scale bar: 200μm

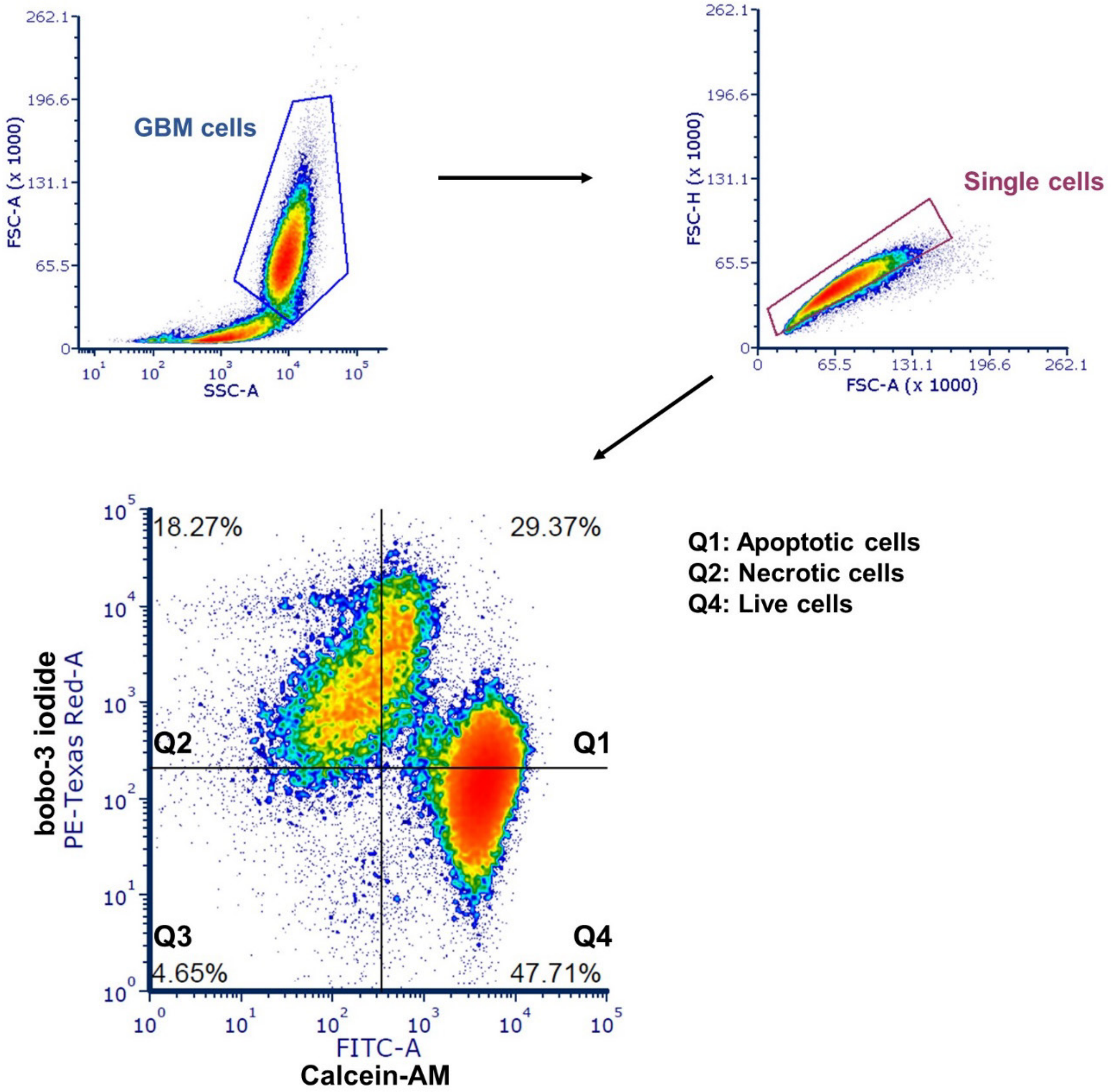


**Figure S2. Cell analysis strategy.** Targets and gating strategy to separate necrotic cells, apoptotic cells, and live cells via flow cytometry using CalceinAM (FITC-A) and Bobo-3 iodide (PE-TexasRed): Live cells (CalceinAM+), apoptotic (CalceinAM+/Bobo-3 iodide+), necrotic (CalceinAM-/Bobo-3 iodide+).

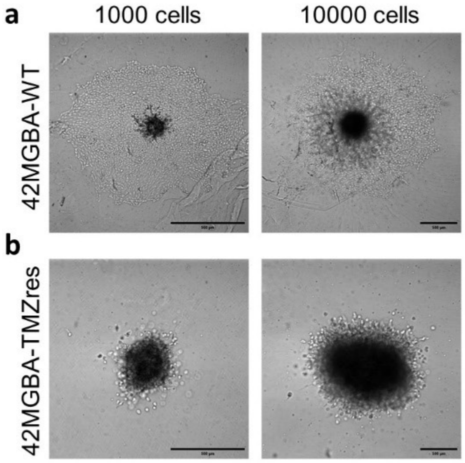


**Figure S3. 42MGBA spheroids treated with DMSO for 5 days after encapsulation.** *(a)* Representative image of 42MGBA-WT spheroids initiated with 1,000 cells (right) or 10,000 cells (left) 5 days after encapsulation and treated with 0.05% DMSO. (b) Representative image of 42MGBA-TMZres spheroids initiated with 1,000 cells (right) or 10,000 cells (left) 5 days after encapsulation and treated with 0.05% DMSO. Scale bar: 500μm.

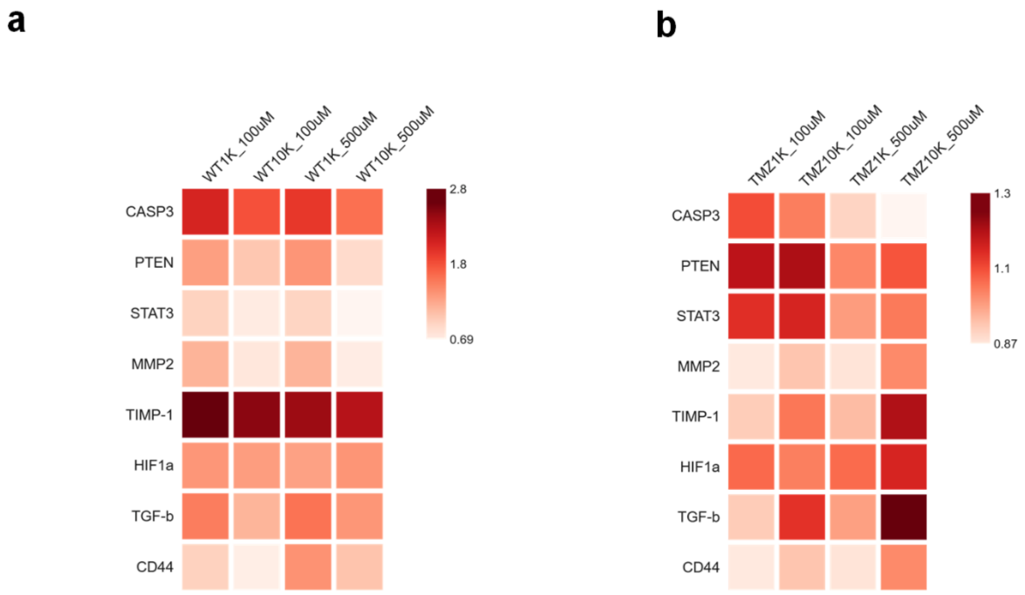


**Figure S4. Shift in gene expression as a function of GBM spheroid size, TMZ treatment, and TMZ-resistance status.** Heatmap of relative gene expression of (a) wild type and (b) TMZ resistant spheroids of different sizes (1,000 vs. 10,000 cells) subject to a single 500 μM TMZ dose vs. 5 days of 100 μM metronomic treatment.
